# Biochemical analysis of deacetylase activity of rice sirtuin OsSRT1, a class IV member in plants

**DOI:** 10.1101/2021.11.05.467508

**Authors:** Nilabhra Mitra, Sanghamitra Dey

## Abstract

The role of plant sirtuins is slowly unwinding. There are only reports of H3K9Ac deacetylation by OsSRT1. This belongs to class IV sirtuin family with a longer C-terminus. Here C-terminus is required for ligand binding and catalysis. OsSRT1 can deacetylate the lys residues at the N terminal tail of both H3 and H4. It can also target the non-histone target, OsPARP1 playing a role in DNA damage repair pathway. Changes in the extent of different histone deacetylation by OsSRT1 is also related with different abiotic stress conditions. NAM and ADP-ribose has negative effect on OsSRT1 deacetylation.

**Highlights:**

1. OsSRT1 is capable of deacetylating various lysine residues of histone H3 and H4 as well as OsPARP1.
2. The extra long C-terminus of OsSRT1 is required for its substrate binding and thus, its catalysis.
3. On plant’s exposure to H_2_O_2_ and Arsenic toxicity, there is a relationship between increased expression of OsSRT1 and increased deacetylation of H3 and H4.

## Introduction

Sirtuins are NAD^+^ dependent deacetylases, divided into five subclasses: I-IV and U. Among all the eukaryotes, human sirtuins are the most studied proteins. Out of the seven human sirtuins, SIRT1 to 3 belong to class I, SIRT4 and 5 belong to class II and III, SIRT6 and SIRT7 belong to class IV [1,2]. SIRT1, 6 and 7 are nuclear/nucleolar, SIRT3, 4 and 5 are mitochondrial and SIRT2 is mainly a cytosolic protein. The two members, SIRT6 and 7 share an overall sequence identity of approximately 38% amongst them.

H3K9Ac and H3K56Ac are the substrates for SIRT6 deacetylation [3–5] but SIRT7 is very specific for H3K18Ac as a lysine deacetylase target [3]. Three-dimensional structures of different human sirtuin constructs are available in apo or/and different ligand complex forms. Sirtuins have a phylogenetically conserved catalytic core and variable N and C terminal regions with respect to length and sequence. The terminal regions of sirtuin proteins have varied functions. The N and C terminus of yeast Hst2 are involved in homotrimer formation, which is responsible for the autoregulation of its activity [4]. Among nuclear proteins, SIRT1 has extensions which regulate its deacetylase activity [5]. The N-terminal end of both SIRT6 and 7 is involved in their enzyme activity and C-terminus for subcellular localization [6,7].

In case of plants, there is limited information available regarding sirtuins’ structure and its related functions. Based on sequence alignment, there are three sirtuins, OsSRT1(OsI_15145), OsSRT2 (OsI_37683) and OscobB (OsI_09569) present in *Oryza sativa indica.* OsSRT1, a class IV member present in nucleus [8], OsSRT2, a classII member in mitochondria [9] and OscobB, a new found class III member. Most of the plants, monocots and dicots, have these two members(SRT1 and SRT2), except maize having only one sirtuins, SRT1. This class III member is also present in few other plants, including rice. OsSRT1(483aa) has homology with both HsSIRT6 (355aa) and HsSIRT7(410aa), thus, belonging to class IV sirtuin family. Based on its nuclear location, the function of OsSRT1 in plant cell is getting unraveled. OsSRT1 is instrumental in deacetylating H3K9Ac at the promoter site of the genes, which are linked with several metabolic processes in plants [8]. It has negative effect on leaf senescence by repressing the genes of methanol-jasmonate cascade [10]. OsSRT1 also represses glycolysis by deacetylating the OsGAPDH and reduces its stress induced accumulation in nucleus [11].

This study will unravel the mysteries of plant sirtuins: How they are linked to PTM in plants and how the PTMs can regulate their action. How strong is their activity and substrate selectivity in plants? The rice sirtuins are present in different cellular locations, nucleus and mitochondria, so it is assumed that they will have varied functions in plants. Sirtuins are known to require NAD^+^ for their activities, which we will confirm in this study. They may affect the level of NAD^+^ and thus the other pathways in the cells. An important question can be answered in this study: Do they have any role in NAD^+^ metabolism in plants as NAD^+^ plays an important part as signal molecule in stress response and pathogenesis in plants. RNAi studies show that sirtuins are involved in stress response in rice plant, especially, under oxidative stress, identification of different modulators will be of great help in this direction. If we can understand the mechanism of sirtuin action under these conditions and detect the interacting partners regulating sirtuins, we can utilize this knowledge to design modulators for sirtuins to make the plants’ more tolerant to stress conditions.

In this work, we are highlighting the acetylated lysine residues present in histone H3 and H4, which are targets of OsSRT1 deacetylation process. Further we have tried to correlate these deacetylation events with different stress conditions. We also noticed that OsSRT1 is capable of influencing the DNA repair pathway as the constituent proteins involved in this pathway can interact with OsSRT1 and eventually get deacetylated. We could identify the role of flanking N and C terminus with respect to substrate specificity and catalysis. In addition, regulation of this deacetylation has been discussed.

## Material and Methods

### 2.1 Reagents

Restriction enzymes, Taq DNA polymerase and T4 DNA ligase were from New England Biolabs, USA. Primers were synthesized from IDT DNA, USA. All the chemicals of analytical grade were procured from Himedia, SRL chemicals and Sigma Aldrich Co., St. Louis, MO. Anti His, anti H3, anti H4 (ab177840), anti PARP1 (ab110915), rPARP1(ab123834), Acetylated H4 antibody pack (ab218056), Acetylated H3 antibody pack (9927T), NHEJ antibody (ab213658) were purchased from Abcam, USA. H3K9Ac peptide (ARTKQTARK(Ac)STGG) was synthetically prepared from Bio Basic Inc., Canada. Anti OsSRT1 against HIS-GST tagged core protein was custom synthesized from rabbit (Biobharati Pvt Ltd, India).

### 2.2 Cloning, overexpression and purification of OsSRT1 constructs

The mRNA was extracted from the rice leaves using the kit (Himedia-HiPurA™ Plant and Fungal RNA Miniprep Purification Kit). This sample was converted into cDNA using Verso cDNA synthesis kit (Thermo Scientific, USA). The *OsSRT1* gene was pcr amplified from the resultant cDNA using the gene specific primers. The *OsSRT1*gene was inserted in pET30a expression vector between *BamHI and EcoRI* restriction sites. The H134Y mutant was prepared using site directed mutagenesis protocol. All the DNA sequences of the OsSRT1 constructs were checked by a DNA sequencing service.

All the constructs were overexpressed and purified following a previous protocol [12]. For oligomerisation analysis of His-OsSRT1 (FL), the purified protein was loaded on an analytical size exclusion column (SEC, superdex200 10/300, GE Healthcare, USA) pre-equilibrated with buffer containing 25mM Tris (pH 8.0), 10% glycerol, 5mM DTT at a rate of 1ml/min (run by AKTA protein purification system). The purified proteins were run on 12 % SDS-PAGE and found to be more than 90% pure. The final yield of OsSRT1 and its other constructs were within the range of 0.8mg per gram of cell. ΔN1 and ΔC1 has low yield (0.2mg/ml).

### 2.3 Interaction studies of OsSRT1 constructs with histones and PARP1(50ug)

Ni pull down experiments were performed using 3μl of His tagged OsSRT1 constructs (0.3μg/ul) and 1μg recombinant human Histone H3 (NEB) and mixed with 20 μl of Ni-NTA slurry (Qiagen). The reaction mix slurry was incubated at 4°C for 4hrs. Then it was centrifuged at 5000g, 4 °C for 10 minutes. The flowthrough was collected and the beads were thoroughly washed several times with buffer (50 mM Tris buffer pH 8.0, 10 mM NaCl, 10% glycerol, 1mM PMSF and 1mM DTT) and finally centrifuged for 10 mins each. The beads were then mixed with a sample dye and boiled. The samples were then resolved using SDS-PAGE. Western blot was performed to study the interaction of different OsSRT1 construct with histone H3. The presence of H3 was detected using anti H3 antibody. Similar experiments were performed to study the interaction of different OsSRT1 constructs and recombinant PARP1 on SDS PAGE using anti PARP1 antibody.

### 2.4 *In vitro* lysine deacetylation activity assay

The deacetylase reaction of purified OsSRT1 was performed at 37°C for 1 hour. Buffer composition for the reaction was 50mM Tris pH 7.5, 150mM NaCl, 5mM DTT and 1mM NAD^+^, 10% glycerol. A 40μl of reaction mixture was set containing 1μg OsSRT1 and mutants, 1μg H3 and volume was made up by buffer. The reaction was stopped by adding the sample dye and boiling it. The reaction mix was then resolved using 12% SDS-PAGE and immunoblot analysis was done by initially blocking the membrane in 5% milk + TBS for 20mins. HRP conjugated rabbit monoclonal H3K18Ac antibody was used to detect the deacetylase activity of the different OsSRT1 constructs. Saturation kinetics for deacetylation were performed with 0-600 μM [NAD^+^] with a fixed concentration of histone H3 (60 nM). Reaction mixtures were incubated at 37 °C for 30 min and contained 5 nM OsSRT1 enzyme. All the experiments were conducted in triplicates.

### Dot Blot Analysis of OsSRT1 deacetylase activity

The reaction mixtures were prepared and carried out as mentioned above. The finished reactions were desalted to remove the unused Bio-NAD^+^ or other buffer constituents. The reaction mix was dot blotted on a nitrocellulose membrane and air dried. The blots were developed as in western blot protocol. The standard curve of both Bio-NAD^+^ and NAD^+^ were prepared for the respective reactions. The intensity of the dots was analyzed using ImageJ software (NCBI) and excel. Saturation kinetics using Non-Linear regression (Michaelis Menten Model) was prepared using GraphPad Prism version 8.3.0 (San Diego, USA).

## Results and discussion

### 1. Analysis of Deacetylase activity of OsSRT1

Ni pull down assays of OsSRT1with the nuclear extract of rice leaves showed that this enzyme can directly interact with both histones, H3 and H4 [12]. We also found that OsSRT1 can deacetylate both these histones. **Fig 1A** shows a timeline of H3 deacetylation by OsSRT1. Literature studies have mostly shown OsSRT1 to deacetylate the acetylated Lys^9^ of histone H3. So, the question was to determine whether OsSRT1 can deacetylate any other acetylated lysine positions of histones or is just specific for the acetylated K9 position like its human homolog SIRT7. HsSIRT7 is absolutely specific for only H3K18Ac [3]. The *K_m_* of NAD^+^ for H3K9Ac deacetylation was found to be 168±25 μM. The *K_m_* and *V*_max_ of the H3K9Ac peptide is 88 ± 8 μM and 5.1 ± 0.15 min^-1^. The concentration of NAD^+^ is kept constant at 250 μM in this reaction. To answer the above question, western blot analyses were performed using the site-specific acetyl antibodies of H3 and H4. Besides H3K9Ac, this enzyme can also deacetylate the acetyl group present at Lys^18^ and Lys^56^ positions of H3 (**Fig 1B and 1C**). OsSRT1 is also capable of deacetylating the acetyl group present at Lys^5^, Lys^12^ and Lys^16^ positions of histone H4 **(Fig 2)**. Interestingly, HsSIRT7 cannot deacetylate H4 whereas HsSIRT6 shows H4 deacetylation, only in presence of nucleosomes [13].

**1).**
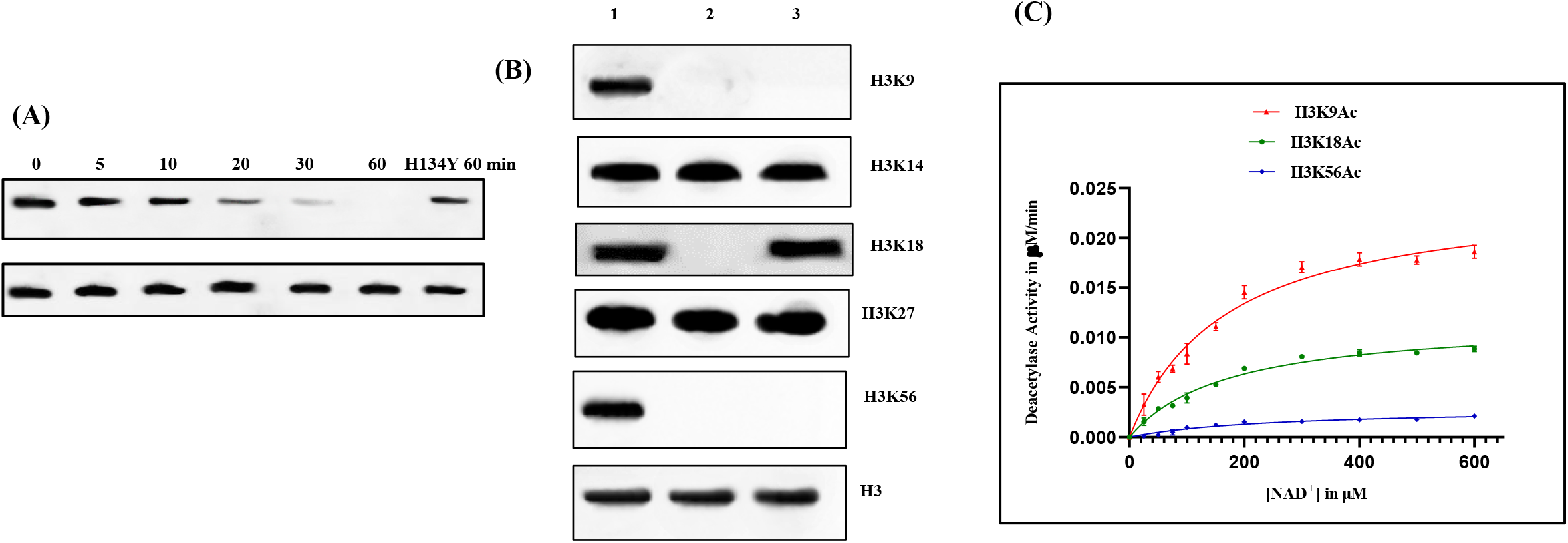
Histone (H3) Deacetylation: (A) Timeline of H3 deacetylation by OsSRT1. (B) **Detection of different acetylated lysine position in H3 which gets deacetylated by OsSRT1:** The deacetylation reaction of OsSRT1 was conducted using nuclear extract prepared from rice leaves at 37°C for 1 hour. In the western blot analysis using site specific antibody we see that OsSRT1 is capable of deacetylating the acetylated K^9^, K^18^ and K^56^ residues no acetyl bands are visible in those lanes. Lane 1 contains empty vector as the negative control, lane 2 contains OsSRT1 in the reaction mix and lane 3 contains HsSIRT6. The lowest panel shows a representative loading control as Histone H3 from nuclear extract. (C) **Saturation kinetics:** The MM plots of OsSRT1 for Histone H3 deacetylations were calculated using Graphpad Prism 9.1.2 (MM Model). The error bar depicts the S.D.; n=3.

**2).**
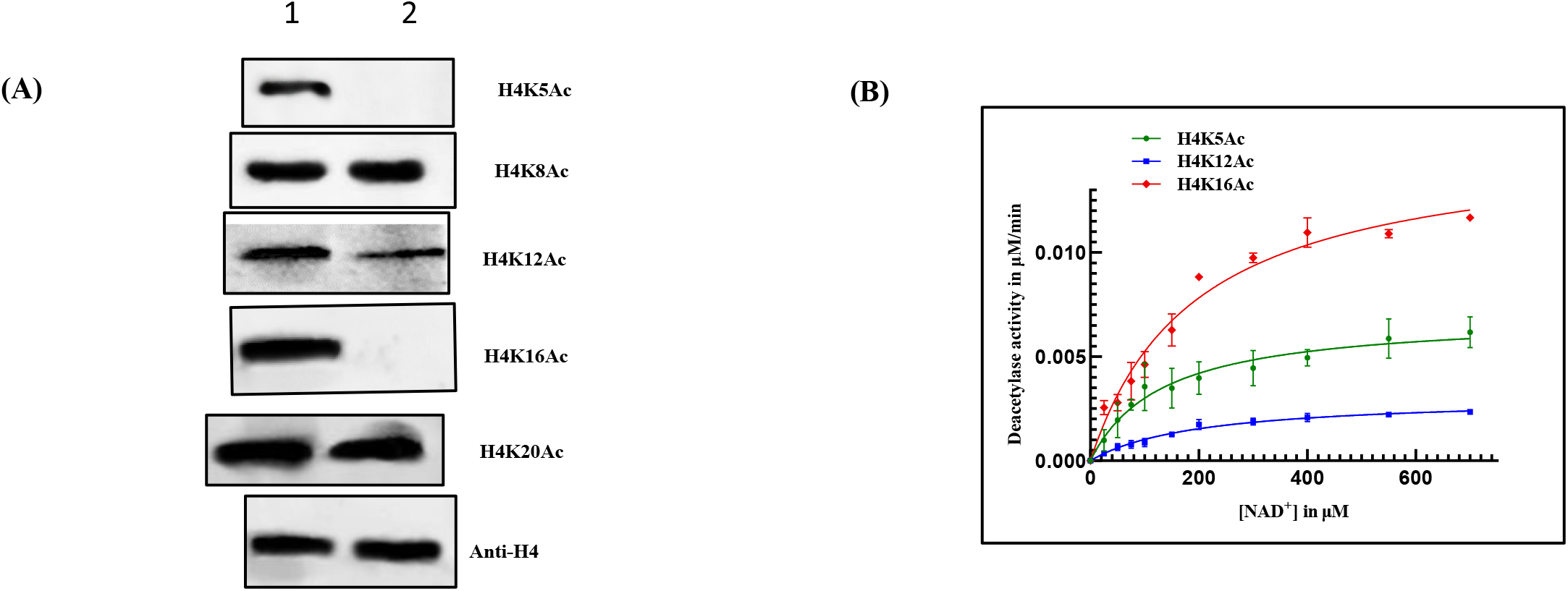
H4 Deacetylation by OsSRT1: **(A) Detection of different acetylated lysine position in H4 which gets deacetylated:** The deacetylation reaction of OsSRT1 was conducted using nuclear extract prepared from rice leaves at 37°C for 1 hour. In this immunoblot, it is seen that OsSRT1 is capable of deacetylating H4K5Ac, H4K12Ac and H4K16Ac. The lowest panel shows a representative loading control as Histone H4 from nuclear extract. Lane1 represents empty vector and Lane 2 contains OsSRT1. (B) **Saturation kinetics:** The MM plots of OsSRT1 for Histone H4 deacetylations were calculated using Graphpad Prism 9.1.2 (MM Model). The error bar depicts the S.D.; n=3. All the assays were conducted by dot blot method. Here the assay was conducted with varied NAD^+^ concentration (0-600uM) with constant histone (H3/H4) concentration from nuclear extract.

OsSRT1 is not that efficient in deacetylating H3K56Ac as noticed from its catalytic efficiency of 0.14 x 10^6^ M^-1^ s^-1^ in comparison to H3K9Ac (1.76 x 10^6^ M^-1^ s^-1^). From all these observations, it seems that OsSRT1 deacetylation shares a combined selection of histone targets of both its human class IV members in nucleus **(TABLE 1)**. OsSRT1 can only utilize NAD^+^ for this chemical reaction (data not shown). We have tested NADP^+^, thio-NAD^+^ and NADH for their involvement in this reaction. Further, we looked at the regulation of OsSRT1 deacetylation. It requires NAD^+^ for its activity and also Zn^2+^ ion for structural stability. OsSRT1 shows immense product inhibition for its activity. NAM and ADP-ribose have IC_50_ value of 147 ± 21 μM and 25 ± 2 μM for H3K9Ac deacetylation. We also found that resveratrol has positive effect on this deacetylation (EC_50_ = 236 ± 12 μM). TricostatinA, an HDAC inhibitor, did not have any effect on OsSRT1 deacetylation whereas sodium butyrate inhibited this reaction with a IC_50_ value of 98 ± 15 μM. TricostatinA has been found to be effective against HsSIRT6 deacetylation (K_i_=2-5μM), binding in the region of acetyl substrate site [14].

**TABLE 1:** A summary of the Michaelis Menten parameters for the deacetylase activity of the OsSRT1 constructs. Kinetic parameters of NAD+ as a substrate for OsSRT1 deacetylase activity of H3K9Ac was measured using dot blot method. This table also shows the catalytic efficiency of different OsSRT1 constructs suggesting the importance of C terminus. Reactions were carried out with 0–600 μM NAD^+^ with constant concentration of H3 (6 nM). ± indicate S.D; n = 3. ND–Not detected.

| **OsSRT1 constructs** | **$K_m$ for NAD<sup>+</sup>(μM)** | **$k_{cat}$ (min<sup>-1</sup>)** | **$k_{cat}/K_m$ (M<sup>-1</sup>s<sup>-1</sup>)** |
| --- | --- | --- | --- |
| OsSRT1 FL | 168 ± 14 | 4.9 ± 0.16 | 1.76 x 10<sup>6</sup> |
| OsSRT1_cc | 256 ± 21 | 2.059 ± 0.076 | 0.48 x 10<sup>6</sup> |
| ΔC1 | 299 ± 43 | 1.2 ± 0.084 | 0.24 x 10<sup>6</sup> |
| ΔC2 | N.D. | N.D. | N.D. |
| ΔN1 | 175 ± 27 | 3.7 ± 0.22 | 1.27 x 10<sup>6</sup> |
| ΔN2 | N.D. | N.D. | N.D. |

### 2. Role of N and C terminal regions of OsSRT1

Multisequence alignment of the three rice sirtuin members shows that OsSRT1 has longer C terminus (approx. 232aa) and OsSRT2 has longer N terminus (approx. 100 aa) in comparison to their counterparts **(Fig 3)**. We wanted to understand the possible roles of these extra terminal regions in OsSRT1 with respect to various processes like substrate binding or its catalysis. We figured that these terminals are not required for its oligomerisation as this enzyme exists as a monomer in solution based on the size exclusion data. For further characterisation, six constructs of OsSRT1 protein were prepared: full length (FL: 1-483), catalytic core region (CC: 45-255), N terminus with CC region (ΔC1: 1-255), N-terminus alone (ΔC2:1-45), C terminus with CC region (ΔN1: 45-483), and C-terminus alone (ΔN2: 255-483) **(Fig 4A)**.

**3).**
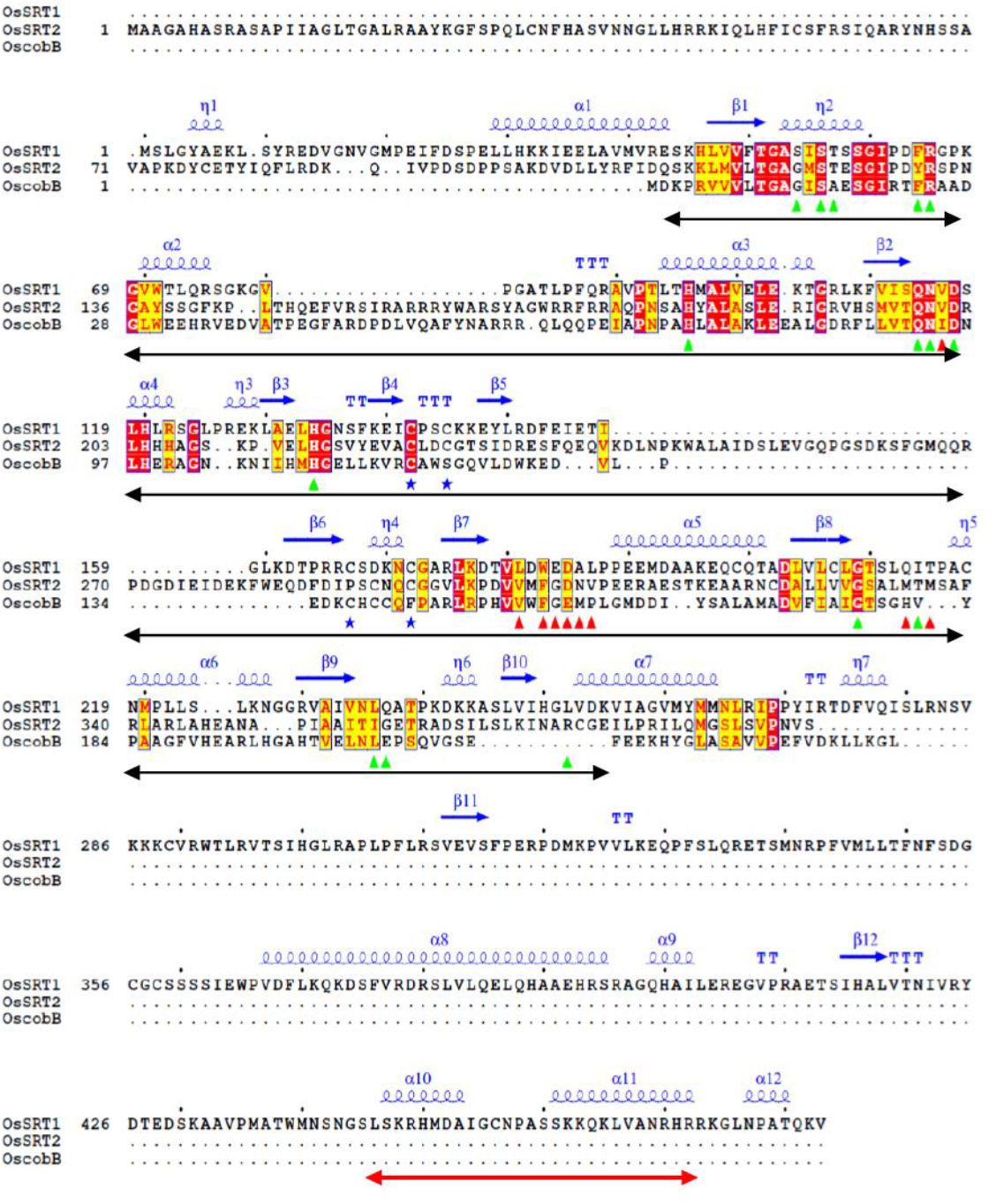
**Sequential analysis of rice sirtuins:** Clustal omega sequence alignment of OsSRT1(B8ARK7) with the other two plant sirtuins OsSRT2 (B8BNG4) and OscobB (A2XBK0). This alignment is showing the major differences with respect to different classes in plants. The residues which line the NAD^+^ binding region are shown in green triangles, peptide binding residues with red triangles, cysteines involved in zinc binding are shown with blue stars. The underlying black arrow demarcates the catalytic core region. The secondary structure depiction is based on the homology model coordinates of OsSRT1. The underlined red arrow at the C-terminus depicts the predicted NLS. This figure is prepared using ESPript 3.0.

We have performed Ni pull down experiments to study the ability of these OsSRT1 constructs to bind the ligands. This study will help us in understanding the involvement of these regions in ligand binding. We observed that full length (FL) and catalytic core (CC) in addition to the ΔN1 construct could bind to both histone H3 and PARP1. ΔC2 bound very weakly with the ligands and showed no activity as it also lacks the catalytic core region. **(Fig 4C)**. Using dot blot analysis, we find comparable activity for FL and ΔC1 construct with weaker activity for CC. Furthermore, CC showed almost two fold lower Km value than the FL for NAD^+^**(Table 1)**. These values suggest the need for the C-terminus in catalysis. Using subcellular prediction software, we detected a region in C terminus involved in nuclear localisation. Moreover, this C terminus is also site for other PTM as analysed by FindMod [15]. Protein Blast analysis of this C-terminal region of *Oryza sativa indica* with other monocots shows up to 50-100 % conservation whereas with dicots it is approx. 40-53% with maximum sequence coverage. This may suggest a similar role of C-terminus in this sirtuins (SRT1) in other plant species.

**4).**
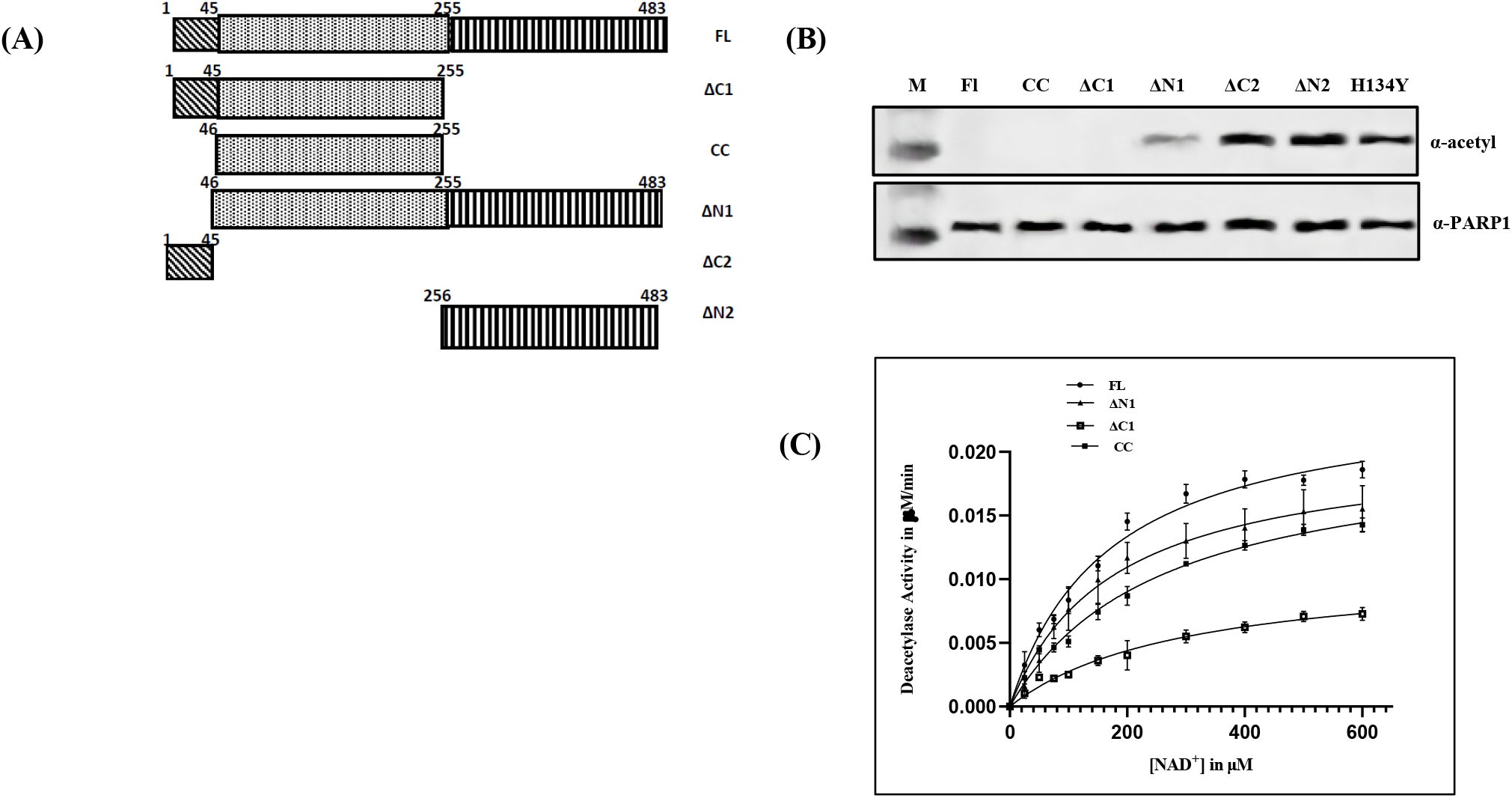
**(A) Schematic diagram of different constructs of OsSRT1:** This figure depicts the demarcation between the different regions of the enzyme. In this study, all the OsSRT1 constructs are prepared based on this distinction. **(B)** Deacetylation of PARP1 by different OsSRT1 constructs. **(C) Saturation enzyme kinetics of H3K9Ac deacetylation by different OsSRT1 constructs:** The MM plots for Histone H3K9Ac deacetylation by different truncates were calculated using Graphpad Prism 9.1.2 (MM Model). The error bar depicts the S.D.; n=3 using dot blot assays.

### 3. Biological role of OsSRT1 deacetylase activity under stress conditions

The role of OsSRT1 in plant is slowly unravelling. As OsSRT1 is localized in the nucleus, we found that it can deacetylate the histones H3 and H4 quite well. As the number of sirtuins are limited in plants it seems that it has combined catalytic abilities of both its human homologs, SIRT6 and SIRT7. Though OsSRT1 can interact with both OsPARP1 and OsPARP2 [12], we found that OsSRT1 can only deacetylate OsPARP1 but not OsPARP2. Ni pull down experiment of OsSRT1 with the nuclear extract of rice leaves has also shown interaction with other DNA damage repair members like H2Ax Ku70, Ku80 and gamma H2Ax. This suggests that OsSRT1 deacetylation might play a role in the DNA damage repair mechanism. Deacetylation by this sirtuin can influence the DNA repair pathway under stress conditions.

There is also a significant increase in deacetylation of specific positions of H3 (K^9^, K^18^, K^56^) and H4 (K^5^, K^12^, K^16^) under abiotic stress conditions. **(Fig 5A & 5B)**. Chromatin modulation in the form of histone deacetylations is also deeply linked to DNA damage repair [16]. It needs to be further seen how the increased deacetylation of different H3 and H4 sites affect the various metabolic pathways in plants under these circumstances. There might be a close relationship between the increased expression of OsSRT1 with the different DNA repair pathway under stress conditions. In addition, both histone ADPr and deacetylation are also linked to DNA repair pathways. Though much is not known about this in plants. Here we are providing some clues in this direction.

**5).**
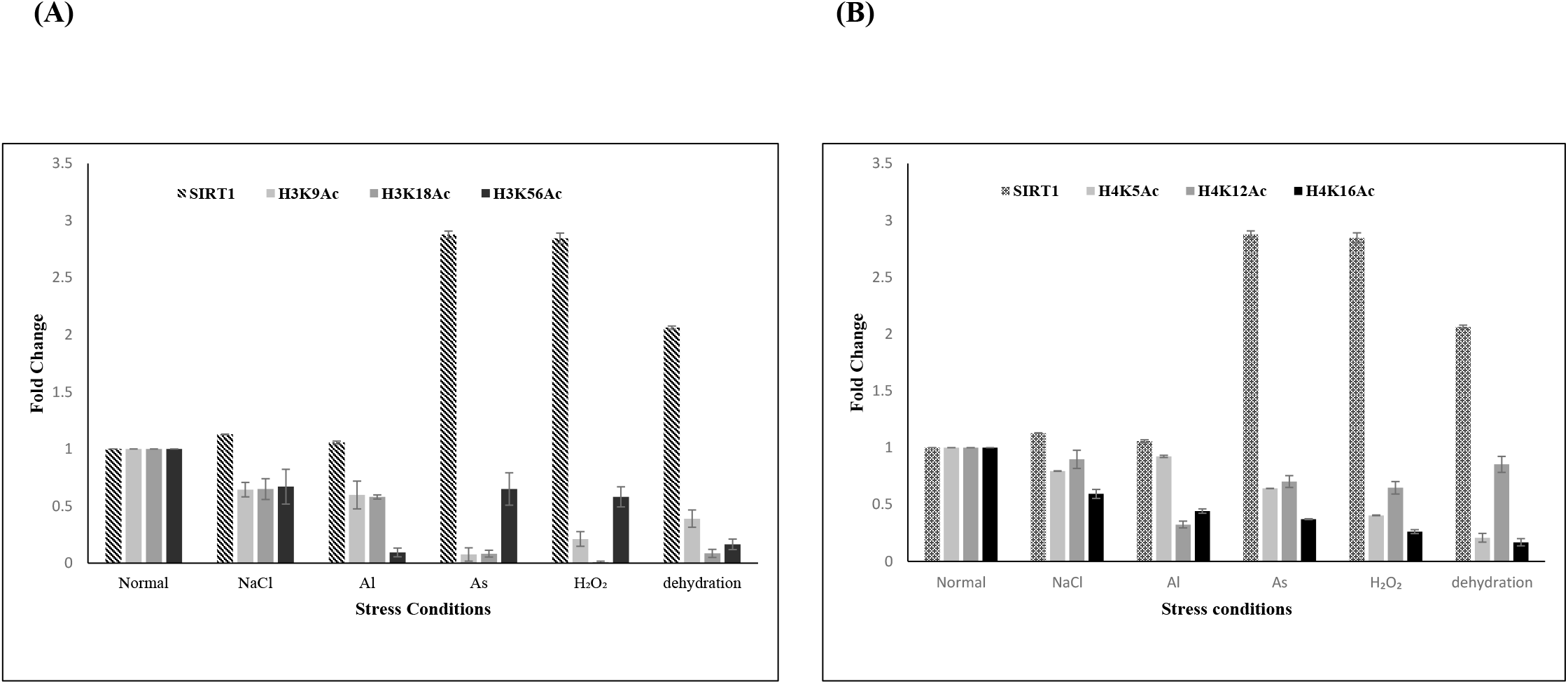
**Link between OsSRT1 upregulation and histone site specific deacetylation**: Western blot analysis of the nuclear extract of rice leaves exposed to varied stress conditions was performed using OsSRT1 and site-specific acetylated lysine antibodies of **(A)** H3 and **(B)** H4 to detect its intensity. The band intensity is measured using ImageJ and the values are normalized before plotting (Microsoft Excel). The histograms suggest a possibility of increase in the deacetylation of H3 and H4 sites by the upregulated OsSRT1 under Arsenic, H_2_O_2_, and dehydration conditions.

## Conclusions

OsSRT1 is almost ubiquitously present in the major tissues of the rice plant. Kinetic studies of its deacetylase activity have been carried out with the revelation of other target sites in histones (most efficient deacetylation with H3K9Ac as substrate followed by H4K16Ac). It needs to be seen which metabolic pathways are affected due to these changes in H3 and H4. The structural features of this plant sirtuin is highlighted with a separate C-terminal domain, which is required for its catalysis. It is seen that the OsSRT1 dual activity are sensitive to the products of the enzyme reaction, NAM and ADPr. NAM can play a major role in balancing the excessive breakdown of NAD^+^ in response to stress conditions, thus maintaining the metabolite homeostasis in cells. Resveratrol, a plant polyphenol has a positive effect on this enzyme action. Thus, there is a possibility of production of transgenic plants overexpressing this molecule, which are more tolerant to the abiotic stress conditions. Further search for more effective activators can be carried out with the screening of resveratrol analogs. Different DNA repair proteins can coordinate amongst themselves to detect and counteract the DNA damages in plants under stress conditions. It needs to be seen how OsSRT1’s dual activity is involved in this whole process.

## Abbreviation

HDAC: Histone deacetylase,
PARP1: Poly ADP-ribose polymerase 1,
PARP2: Poly ADP-ribose polymerase 2

## Acknowledgement

This work was funded by Department of Science and Technology, Govt of India (EMR/2014/000366) and FRPDF scheme, Presidency University, Kolkata, India.

